# Hedgehog signaling is non-cell autonomously activated in the cystic kidney of *Arl13b* mutant mice

**DOI:** 10.1101/2022.02.28.482403

**Authors:** Chia-Ling Hsieh, Stephanie Justine Jerman, Zhaoxia Sun

## Abstract

**Background:** Polycystic kidney disease (PKD) is a ciliopathy characterized by fluid-filled epithelial cysts in the kidney. Although it is well established that the primary cilium is essential for Hedgehog (HH) signalling and HH signalling is abnormally activated in cystic kidneys of multiple PKD models, the mechanism and function of HH activation in PKD pathogenesis remains incompletely understood.

**Methods:** We used a transgenic HH reporter line to identify the target tissue of HH signalling in *Arl13^f/f^;Ksp-Cre* mutant kidney, in which the cilia biogenesis gene *Arl13b* is specifically deleted in epithelial cells of the distal nephron. In addition, we treated *Arl13b^f/f^;Ksp-Cre* mice with the GLI inhibitor GANT61 and analyzed its impact on PKD progression in this model.

**Results:** *In vivo* in the mouse kidney, deletion of *Arl13b* in epithelial cells led to non-cell autonomous activation of the HH pathway in the interstitium. In addition, whole body inhibition of the HH pathway by GANT61 reduced cyst burden, suppressed fibrosis and reduced kidney function decline in *Arl13b^f/f^;Ksp-Cre* mice.

**Conclusions:** Our results reveal non-cell autonomous activation of HH signalling in the interstitium of the cystic kidney of *Arl13b^f/f^;Ksp-Cre* mice and suggest that abnormal activation of the HH pathway contributes to PKD progression.

## INTRODUCTION

The primary cilium is a microtubule-based cellular organelle that protrudes from the cell surface into the extracellular matrix or fluid, and functions as an antenna to sense and transduce environmental signals to regulate cellular responses^1–3^. One of the best studied cilia-regulated pathways is the Hedgehog (HH) pathway. Three secreted ligands, Sonic hedgehog (SHH), Indian hedgehog (IHH) and Desert hedgehog (DHH), initiate HH signaling by binding to the receptor PATCHED1/2 (PTCH1/2). In vertebrates, PATCHED is localized to cilia in the absence of its ligands. Binding of ligands activates the HH pathway by displacing ciliary PTCH and allows the G-protein coupled receptor-like protein SMOOTHENED (SMO) to enter the cilium. SMO accumulation facilitates processing of GLI proteins, which are then trafficked to the nucleus and transcribe HH target genes, including *Gli1, Ptch1* and *Ptch2*^4–6^. A functional cilium is therefore required for the activation of the HH pathway. The HH signaling pathway plays a fundamental role in tissue patterning, cell growth and differentiation during development^7–9^. HH signaling is frequently reactivated during injury and repair in multiple organs, including the lung and kidney, and prolonged HH activation contributes to tissue fibrosis^10–14^.

Mutations in ciliary genes, encoding proteins required for cilia formation or function, contribute to a wide spectrum of human disorders, including polydactyly, holoprosencephaly, obesity and polycystic kidney disease (PKD), among many others, collectively known as ciliopathies^15–17^. While abnormal HH signaling underlies multiple ciliopathies such as polydactyly and craniofacial ciliopathies, the molecular mechanism of PKD pathogenesis remains poorly understood^18,19^. Autosomal dominant polycystic kidney disease (ADPKD), the most frequent form of PKD and one of the most common monogenetic diseases in human, can be attributed to mutations in *PKD1* and *PKD2*, which encode POLYCYSTIN 1 and 2 (PC1 and PC2), respectively (reviewed in ^20^). PC1 and PC2 are trafficked to cilia and their ciliary localization is integral to their function^21–25^. It is postulated that PC1 and PC2 functions to inhibit a cilia-dependent and cyst promoting pathway (CDCA) and unconstrained CDCA leads to cyst formation in *Pkd1* or *Pkd2* mutants^26^. However, the precise molecular mechanism of this pathway is undefined.

Abnormal activation of the HH pathway has been detected in cystic kidney samples from both human and animal models, including *Thm1* mouse mutants^27–30^. *Thm1/Ift139* encodes a component of the IFT A complex that negatively regulates HH signaling^31^. Defective *Thm1* leads to shorter cilia with bulbous tips, upregulation of HH signaling and cystic kidney^27,31^. In concordance with *Thm1’s* inhibitory role in HH signaling, repressing the HH pathway reduced cyst progression in the *Thm1* mutant kidney^27^. Moreover, cyst expansion induced by *cAMP* and *Pkd1* inactivation in cultured mouse kidney explants was also shown to be sensitive to HH inhibition^27^. In the meantime, a recent study demonstrated that modulating the HH pathway in renal epithelial cells failed to affect PKD progression in mouse *Pkd1* mutants^32^, suggesting that the role of HH signaling in different PKD models is complex and not fully understood.

In a previous study, we found that the HH pathway is upregulated in the cystic kidney of *Arl13b^f/f^;Ksp-Cre* mice, where *Arl13b* is specifically deleted in epithelial cells of the distal nephron^29^. Since *Arl13b* is required for cilia biogenesis, this result seems contradictory to the essential role of cilia in HH signaling. However, HH ligands commonly activate HH signaling non-cell autonomously during tissue patterning^8,9^. Similarly, following acute kidney injury, SHH is produced by epithelial cells, but downstream targets are upregulated in interstitial cells^10,12,33^. The role of non-cell autonomous HH signaling has not been investigated in PKD pathogenesis and is the focus of this study.

We demonstrate that HH signaling is predominantly activated in the interstitium when *Arl13b* is deleted in renal epithelial cells. In addition, treatment of the GLI inhibitor GANT61 partially suppressed both renal cyst expansion and fibrosis, and preserved kidney function in *Arl13b^f/f^;Ksp-Cre* mice. Together, our results clarify the role of renal epithelial and interstitial cells in the abnormal activation of HH signaling in cystic kidneys and highlight the significance of epithelial-interstitial cross-talk in this process. Our results also suggest that non-cell autonomous HH signaling contributes to both renal cyst progression and fibrosis in *Arl13b* mutant mice, and may provide important insight into the molecular etiology of cystic diseases in human.

## RESULTS

### Hedgehog signaling is activated non-cell autonomously in the interstitium of the *Arl13b^f/f^;Ksp-Cre* kidney

We previously showed that the expression of HH ligands and target genes are upregulated in the cystic kidney of *Arl13b^f/f^;Ksp-Cre* mice in whole kidney lysates^29^. To ascertain tissue specificity of HH activation in the mutant kidney, we took advantage of the *Gli1*^*LacZ*/+^ reporter line that expresses nuclear localized LacZ in cells with activated HH signaling^34^. We generated *Arl13b^f/f^;Ksp-Cre; Gli1*^*LacZ*/+^ mice and performed immunofluorescence analysis for LacZ in sections of postnatal day 21 (P21) kidneys. In kidneys with functional ARL13B (*Arl13b^f/+^;Ksp-Cre;Gli1^LacZ/+^*), we detected LacZ positive nuclei sporadically in both the cortex and medullar region, outside of epithelial tubules, as outlined by anti-Laminin signal, consistent with the known distribution of *Gli1*^+^ cells in the wild type kidney (Fig. 1A)^10^. In the *Arl13b^f/f^;Ksp-Cre;Gli1^LacZ/+^* mutant kidney, the number of LacZ positive nuclei increased; notably, they were still excluded from regions encircled by Laminin, suggesting that stromal cells were the HH responding cells (Fig. 1A). Since the morphology of tubules, epithelial cells and the pattern of Laminin signal were significantly distorted in the mutant kidney, to validate the identity of nuclear LacZ (nLacZ) positive cells, we additionally performed co-labelling with anti-LacZ and anti-alpha Smooth Muscle Actin (aSMA), a marker of activated myofibroblasts. In both the cortex and medullar region of the mutant kidney, the signal of aSMA increased dramatically, as expected^29^ (Fig. 1B). Moreover, most nLacZ positive cells were also positive for aSMA, suggesting that they were myofibroblasts (Fig. 1B).

**Figure 1.**
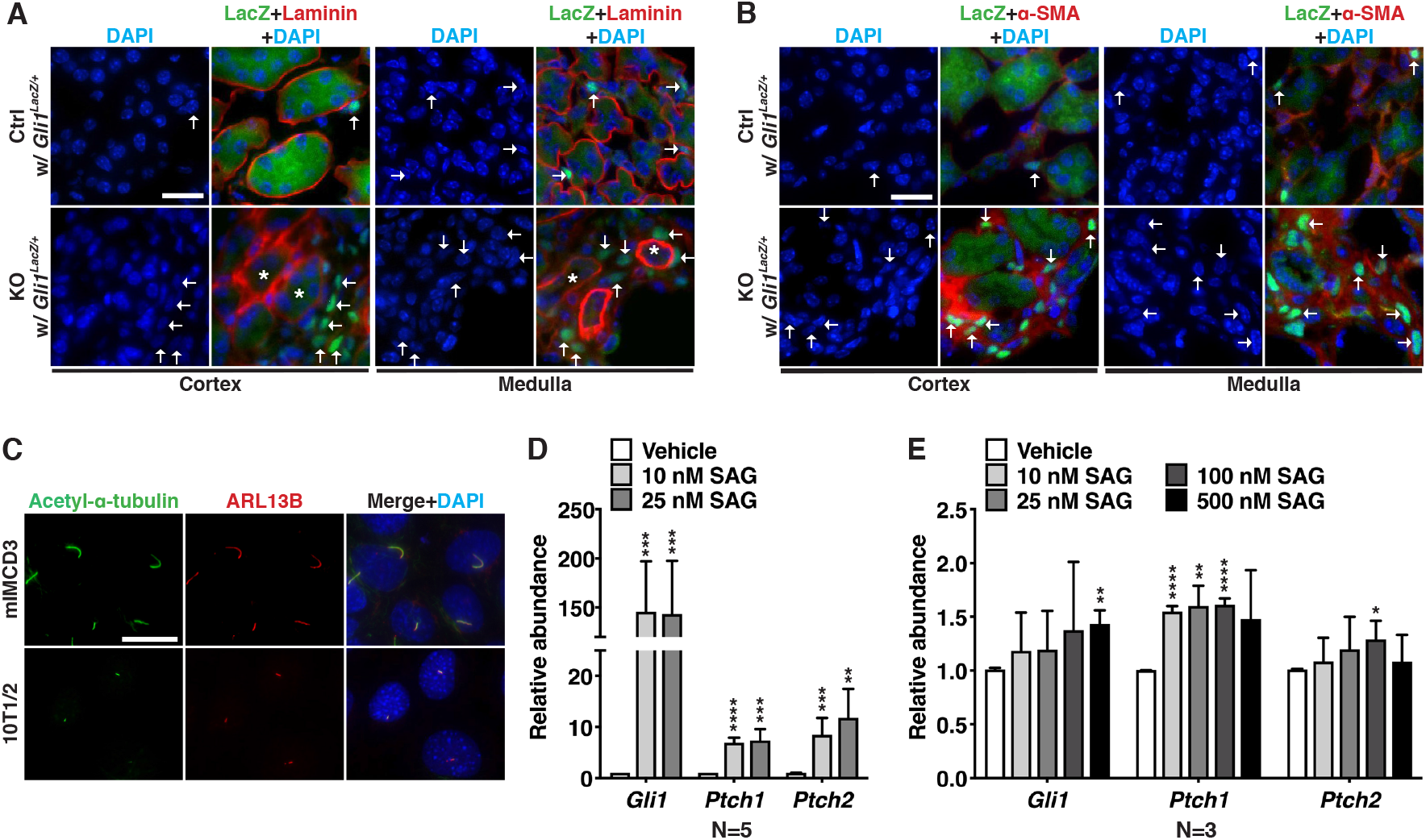
Epithelial-specific inactivation of *Arl13b* results in non-cell autonomous activation of HH signaling in stromal cells. (**A, B**) Immunofluorescent staining of kidney sections of *Arl13b^f/f^;Ksp-Cre;Gli1*^*LacZ*/+^ and *Arl13b*^*f*/+^;*Ksp-Cre;Gli1*^*LacZ*/+^ mice at P21. Cells with active HH signaling are indicated by nLacZ signal (green, detected by anti-LacZ) Arrows point to representative nLacZ positive nuclei. Stars indicate nLacZ negative epithelial region encircled by Laminin. DAPI = nuclear counterstain (blue). Scale bars: 20 μm. In (A), anti-Laminin (red) outlines the basal border of epithelial tubules. In (B), anti-α-SMA (red) labels activated myofibroblasts. (**C**) Cilia in mIMCD3 and 10T1/2 cells detected by anti-acetylated tubulin (green) and anti-ARL13B (red). Nuclei stained with DAPI (blue). Scale bar: 10 μm. (**D, E**) QRT-PCR analysis of the expression level of HH target genes in 10T1/2 (D) and mIMCD3 (E) cells in response to SAG treatment. Unit 1 is defined by the expression level in vehicle treated controls. Results represent mean ± SD of at least three independent experiments. Ctrl: *Arl13b*^*f*/+^;*Ksp-Cre* mice; KO: *Arl13b^f/f^;Ksp-Cre* mice; *P<0.05; **P<0.01; ***P<0.001; ****P<0.0001. All scale bars: 20 μm.

Combined, these results suggest that inactivation of *Arl13b* in renal epithelial cells activates HH signaling in stromal cells via a non-cell autonomous mechanism.

### Mesenchymal 10T1/2 and epithelial mIMCD3 cells show differential responsiveness to HH stimulation

Since epithelial cells in the *Arl13b*^*f*/+^*;Ksp-Cre;Gli1*^*LacZ*/+^ kidney are negative for HH signaling, as indicated by the lack of nLacZ signal (Fig. 1A), we investigated whether mesenchymal and epithelial cells show differential response to HH stimulation. We cultured mouse inner medullary collecting duct (mIMCD3) cells and the murine mesenchymal cell line 10T1/2 *in vitro*. After reaching confluency, both cell lines were switched to a low serum medium (0.5% fetal bovine serum (FBS)) to induce the formation of primary cilia. As expected, abundant cilia could be detected in mIMCD3 cells (Fig. 1C). In addition, 10T1/2 cells also displayed cilia, shown by cilia markers anti-acetylated tubulin and anti-ARL13B (Fig. 1C). Ciliated cells were subsequently treated with SAG, a small molecule agonist of SMO, for 24 hours. The expression level of HH target genes, including *Gli1, Ptch1* and *Ptch2*, was then analyzed by quantitative RT-PCR (qRT-PCR) to monitor the status of HH signaling. Even at a low SAG concentration (10nM), expression levels of HH target genes *Gli1* (>143-fold), *Ptch1* (~7-fold) and *Ptch2* (12-fold) were significantly increased in stromal 10T1/2 cells compared to vehicle treated controls (Fig. 1D). In contrast, and despite that mIMCD3 epithelial cells displayed more prominent cilia (Fig. 1C), mIMCD3 cells showed minimal response to SAG, with a slightly increased expression of *Gli1* (1.4-fold) at a much higher SAG concentration (500 nM), *Ptch1* (1.5 fold at 10 nM SAG) and *Ptch2* (1.3 fold at 100 nM SAG) (Fig. 1E). The limited sensitivity of mIMCD3 cells to activation of HH signaling is consistent with the lack of nLacZ signal in renal epithelial cells (Fig. 1A).

### The GLI inhibitor GANT61 partially suppresses the renal phenotypes of *Arl13b^f/f^;Ksp-Cre* mice

Since HH signaling is activated non-cell autonomously in the *Arl13b^f/f^;Ksp-Cre* kidney, to investigate the functional significance of this abnormal signaling in PKD progression, we used GANT61, a small molecule that interferes with the transcriptional activities of both GLI1 and GLI2^35^, to achieve whole body HH inhibition. Our previous study established that the *Arl13b^f/f^;Ksp-Cre* kidney is non-cystic at P7 and significantly cystic at P14^29^. Because of technical difficulties of intraperitoneal (i.p.) injection into P7 mice, we analyzed key characteristics of the *Arl13b^f/f^;Ksp-Cre* kidney at P10 and results showed that the *Arl13b^f/f^;Ksp-Cre* kidney was slightly cystic at this stage (Fig. 2). The kidney weight to body weight (KWB) ratio, cystic index and blood urea nitrogen (BUN) level were increased as well in both female and male mutant mice (Fig. 2B-G), although at levels much milder than P14 mutants^29^. We therefore started GANT61 treatment from P10. *Arl13b^f/f^;Ksp-Cre* and control mice (*Arl13b^f/+^;Ksp-Cre*) were subjected to daily i.p. injection of GANT61 or vehicle alone(Figure 3A). After GANT61 treatment, qRT-PCR using total kidney lysates verified that this treatment led to reduced expression level of the HH target genes *Gli1, Ptch1* and *Ptch2* in mutant kidneys, and to a lesser degree in control kidneys (Fig. 3B). Both female and male mutant mice treated with GANT61 showed significantly reduced kidney weight to body weight (KBW) ratio, while no difference was seen in control animals (Fig. 3C, D). Taking advantage of variations in phenotypic severity between individual mice, we asked whether the level of *Gli1* expression correlates with KBW ratio by Pearson correlation analysis. Interestingly, while there was no significant correlation between *Gli1* level and KBW ratio in vehicle treated mutants, the correlation coefficient was statistically significant in GANT61 treated mutants (Fig. 3E, F). Cyst progression, as indicated by cyst index, was also inhibited by this treatment in both male and female mice (Fig. 3G, H, I). By contrast, the same GANT61 treatment had minimal effect on kidney size, KBW ratio and cyst formation in control mice, even though the HH pathway was modestly inhibited in the kidney (Fig. 3B, C, D, H, I).

**Figure 2.**
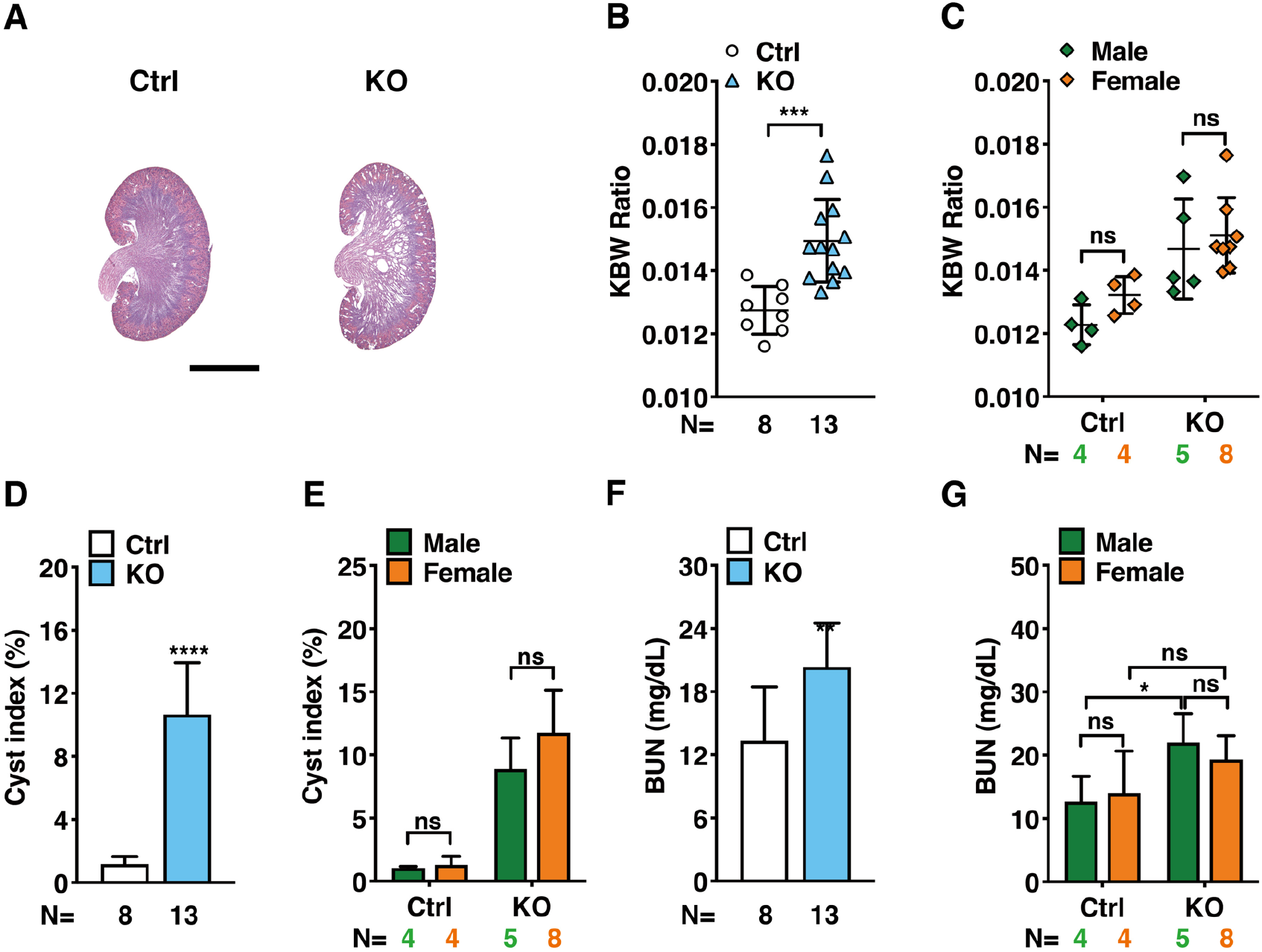
Phenotypes of the *Arl13b*^*f/f*^*;Ksp-Cre* (KO) kidney at P10. *Arl13bf*^*f*/+^;*Ksp-Cre* mice are used as control (Ctrl). Scale bar: 2 mm. (A) Hematoxylin and eosin-stained kidney sections. (B, C) KBW in KO and Ctrl mice. (D, E) Cystic index in KO and Ctrl mice). (F, G) BUN level in KO and Ctrl mice. *P<0.05; **P<0/01; ****P<0.0001; ns, P>0.05.

**Figure 3.**
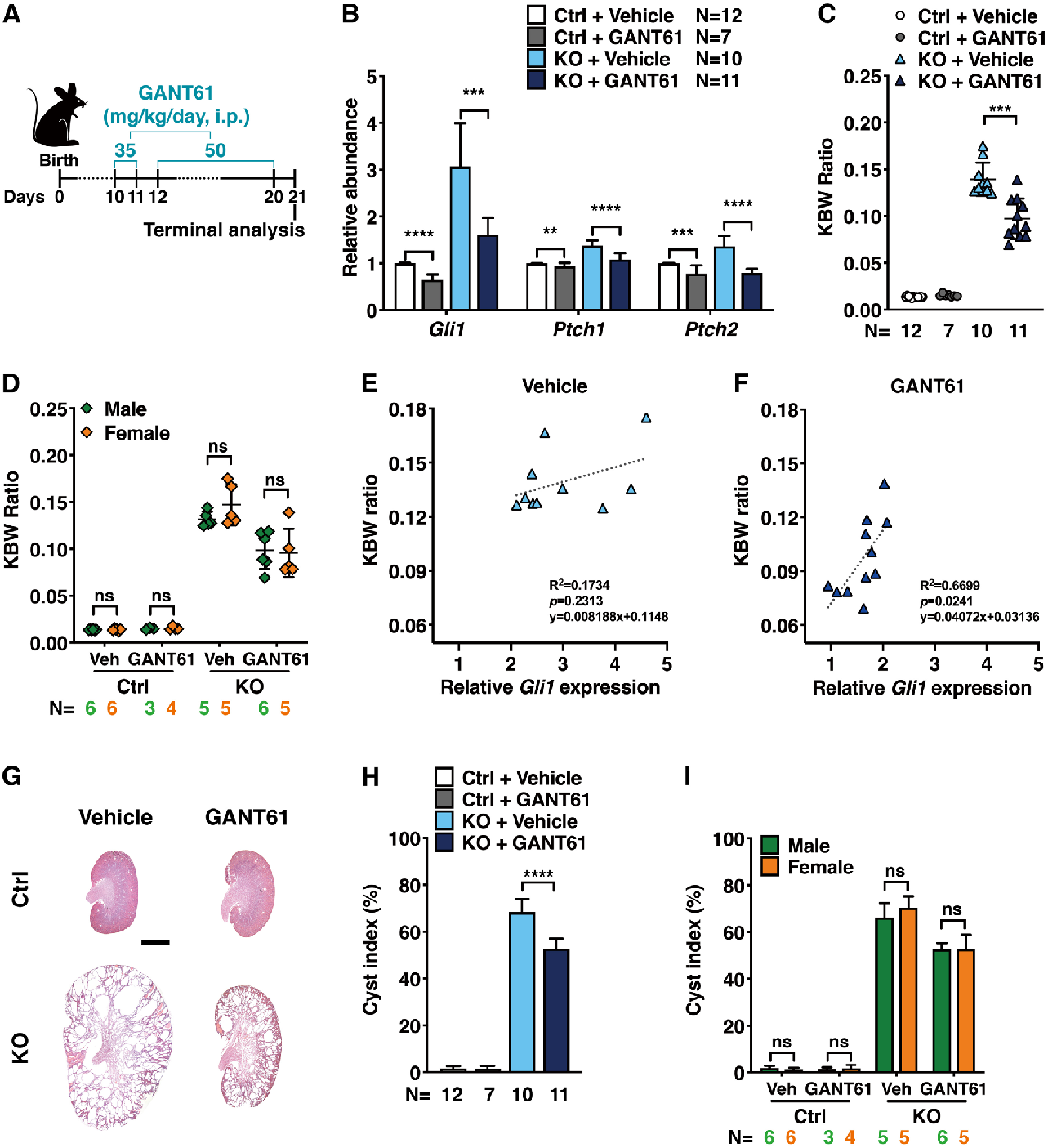
GANT61 suppresses kidney cyst progression in *Arl13b^f/f^;Ksp-Cre* (KO) mice. (**A**) Schematic diagram of the treatment schedule. P10 and P11 mice were subjected to daily i.p. injection of 35mg/kg/day GANT61, followed by daily i.p. injection of 50 mg/kg/day GANT61 from P12 to P20. (**B**) GANT61 treatment reduced the expression level of HH target genes *Gli1, Ptch1* and *Ptch2* in whole kidney lysates of KO and *Arl13b*^*f*/+^;*Ksp-Cre* (Ctrl) mice, assayed by qRT-PCR. *Gapdh* was used for normalization. Unit 1 is defined by the expression level in vehicle treated animals. Results represent mean ± SD of at least three independent experiments. (**C, D**) KWB ratio in vehicle (Veh) and GANT61 treated mutant KO and *Arl13b*^*f*/+^;*Ksp-Cre* (Ctrl) mice. (**E, F**) Pearson correlation between the level of whole kidney *Gli1* expression and KWB in vehicle (E) and GANT61 (F) treated *Arl13b*^*f/f*^;*Ksp-Cre* mice. (**F**) Hematoxylin and eosin-stained kidney sections from GANT61 and vehicle *Arl13b*^*f*/+^;*Ksp-Cre* (Ctrl) and KO mice. Scale bar: 2 mm. (**G, I**) Cystic index of GANT61 and vehicle (Veh) treated *Arl13b*^*f*/+^*;Ksp-Cre* (Ctrl) and KO kidneys. **P<0.01; ***P<0.001; ****P<0.0001; ns, P>0.05.

We then investigated the impact of GANT61 treatment on myofibroblast activation in the kidney of *Arl13b^f/f^;Ksp-Cre* mice. Immunofluorescence analysis of kidney sections revealed that VIM and aSMA signals in the interstitium of the renal cortex and medullar regions were significantly decreased in GANT61 treated mutants (Fig. 4A). We further analyzed collagen deposition by trichrome staining. Results revealed that the blue staining, i.e. collagen deposition, was reduced in treated mutants (Fig. 4B). Moreover, Western blot using whole kidney lysates from mutant mice revealed a notable reduction of the protein level of α-SMA and collagen I by GANT61 treatment (Fig. 4C).

**Figure 4.**
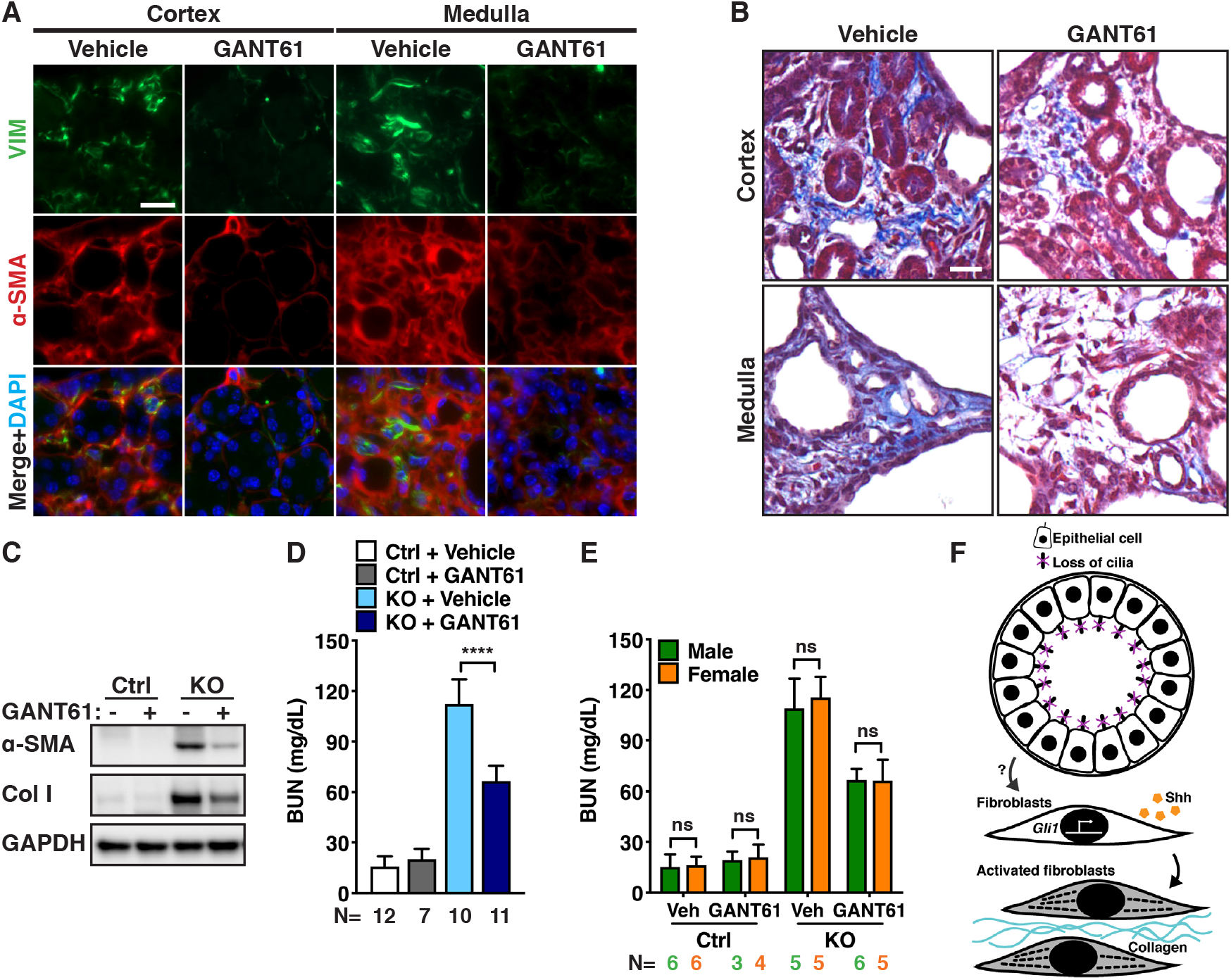
GANT61 suppresses renal fibrosis and improves kidney function in *Arl13b^f/f^;Ksp-Cre* (KO) mice. (A) Immunofluorescent staining of kidney sections of vehicle-and GANT61-treated KO mice. Anti-VIM in green, anti-αSMA in red. DAPI, blue. Scale bar: 20 μm. (B) Trichrome-stained kidney sections of vehicle- and GANT61-treated KO mice. Blue color indicates collagen deposition. Scale bar: 20 μm. (C) Western blot showing the protein level of αSMA and collagen I (Col I) in whole kidney lysates of KO mice treated with GANT61 (+) or vehicle (-). GAPDH is used as a loading control. (D, E) BUN level in GANT61 or vehicle treated KO and *Arl13b*^f/+^;*Ksp-Cre* (Ctrl) mice. Male and female combined is shown in D and separately in E. Results represent the mean ± SD of at least three independent experiments. ****P<0.0001; ns, P>0.05. (F) A model showing the non-cell autonomous activation of HH signaling in stromal cells.

In concordance with morphologic improvement, the BUN level was significantly reduced in GANT61 treated mutants, indicating preserved kidney function (Fig. 4D, E).

Combined, these results show that the GLI inhibitor GANT61 partially suppresses both cyst progression and fibrosis in the *Arl13b^f/f^;Ksp-Cre* kidney.

## DISCUSSION

The primary cilium plays a key role in both PKD pathogenesis and HH signaling. However, the relationship between the two is less clear and results from different studies are contradictory. For example: It was shown that inhibiting HH signaling in mouse mutants of *Thm1*, a negative regulator of HH signaling, partially suppresses cyst progression^27^. However, in mouse *Pkd1* mutant models, cell autonomous inhibition of HH signaling, through genetic inactivation of *Smo* in epithelial cells, does not alter PKD progression^32^. These interesting studies suggest that different PKD or ciliary genes might have distinct relationships with HH signaling, hence the different response to HH inhibition. Alternatively, HH signaling could be activated non-cell autonomously in cystic kidneys, resulting in the lack of response by cell autonomous inhibition of HH. We, therefore, tested the mode of HH activity in a model of cilia dysfunction and PKD, using a HH reporter in *Arl13b^f/f^;Ksp-Cre* mutant mice. We show that when *Arl13b* is deleted in epithelial cells, HH signaling is activated non-cell autonomously in interstitial cells, consistent with the pattern of HH activation during development and tissue repair ^8–10,12,33^. Non-cell autonomous HH signaling could be activated through secreted ligands or through mechanical stress exerted by expanding cysts and activate myofibroblast in the stroma (Fig. 4F).

The severe phenotypic consequences of defective epithelial cilia in the *Arl13b^f/f^;Ksp- Cre* kidney, coupled with the lack of strong HH response in epithelial cells in this model, suggest the existence of a separate cilia-mediated pathway in epithelial cells, disruption of which triggers a cascade of events leading to the formation of epithelial cysts and non-cell autonomous activation of HH signaling in interstitial cells. The relationship between this undefined ciliary pathway and the CDCA pathway, which also functions in epithelial cilia, is unknown. It will be interesting to investigate whether the CDCA pathway plays a role in non-cell autonomous activation of the HH pathway in cystic kidney models. Alternatively, Arl13b could function through an extraciliary pathway^36^. Whether this pathway is active in renal epithelial cells remains to be investigated.

Given the non-cell autonomous nature of HH activation in the *Arl13b^f/f^;Ksp-Cre* kidney, we sought to investigate the role of abnormal activation of HH signaling by using the small molecule GLI inhibitor GANT61 to reduce HH signaling globally. GANT61 treatment attenuated both renal fibrosis and cyst progression in *Arl13b^f/f^;Ksp-Cre* mutants. The impact on renal fibrosis is consistent with the role of HH in renal fibrosis triggered by kidney injury. Previous studies showed that HH signaling is activated after kidney injury and this activation is integral to repair^10,12,33,37,38^. Functionally, both global pharmacological inhibition of HH signaling and targeted genetic deletion of *Shh* in renal epithelial cells attenuated fibroblast activation after kidney injury ^12,33,37,38^. GANT61 could similarly attenuate fibrosis in the *Arl13b^f/f^;Ksp-Cre* kidney through inhibiting fibroblast activation. The reduced level of aSMA and VIMENTIN in treated mutants is consistent with this model. Interestingly, GANT61 treatment also ameliorated cyst progression. Whether this effect is through reduced fibroblast activation or a more indirect mechanism such as macrophage recruitment, which has an established role in cyst progression, remains to be investigated^39,40^. Further, it will be vital for our understanding of PKD to test whether interstitial cells are also HH responsive cells in additional PKD models, including Polycystin mutants and mutants affecting different aspects of cilia biogenesis and signaling.

## METHODS

### Ethics statement

Animal experiments in this study were carried out at Yale University School of Medicine in accordance with the Animal Use Protocols as approved by the Institutional Animal Care and Use Committee.

### Animals

*Arl13b^f/f^;Ksp-Cre* mice have previously described^29^. *Gli1^lacZ^* mice (Jackson Laboratory, Bar Harbor, ME) were provided by the Liem lab. To generate *Arl13b^f/f^;Ksp-Cre;Gl1i*^*lacZ*/+^ mice and control littermates (*Arl13b*^*f*/+^;*Ksp-Cre;Gli1*^*lacZ*/^+), *Arl13b^f/f^;Gli1*^*lacZ*/+^ mice were crossed to *Arl13b*^*f*^/+;*Ksp-Cre* mice.

### GANT61 treatment

GANT61 (ApexBio Technology, A1615, 25mg) was dissolved in ethanol (1.75 ml) and stored at −80°C. For mouse injection, the ethanol solution was further diluted in PBS (3:7) immediately before injection. GANT61 was intraperitoneally administered to male and female mice at a dose of 30 mg/kg body weight (BW)/day at both 10 and 11 days of age, and 50 mg/kg BW/day at 12 to 20 days of age (Figure 4A). Control mice were injected with vehicle alone following the same schedule. Mice were sacrificed at P21.

### Cell culture

mIMCD3 cells were cultured in Dulbecco’s modified Eagle’s medium/Nutrient Mixture F-12 (Invitrogen, 11330-032) supplemented with 10% heat-inactivated fetal bovine serum (FBS, Invitrogen, 16410) and 1% Antibiotic-Antimycotic solution (Gibco, 15240062). C3H/10T1/2, clone8 (10T1/2) cells were obtained from the American Type Culture Collection (ATCC, CCL-226) and was cultured in Basal Medium Eagle (BEM, Gibco, 21010046) supplemented with 10% heat-inactivated FBS, 2mM L-glutamine (Gibco, 25030081) and 1% Antibiotic-Antimycotic solution. Cells were maintained as a monolayer at sub-confluent densities in a 5% CO_2_ humidified incubator at 37 °C. For immunofluorescence analysis, cells were seed into chambered cell culture slide (Fisher Scientific, 08-774-26). To induce primary cilium formation, cells were treated in low-serum (0.5% FBS) medium after reaching confluency and maintained for the duration indicated in each experiment. For Smoothened Agonist (SAG) treatment, SAG (Cayman Chemical, 11914) was dissolved in DMSO to make the stock solution (5mM). After one day of culturing in low serum medium (0.5% FBS), cells were treated with varying concentrations of SAG in medium containing 0.5% FBS for 24 hours or 4 days as indicated in each experiment. The same amount of DMSO was used as vehicle control.

### Western blot using cultured cells and mouse kidney tissues

Cell pellets and kidney tissue samples were homogenized in whole cell extracts (WCE) buffer containing 20 mM HEPES, pH 7.4, 0.2 M NaCl, 0.5% Triton X-100, 5% glycerol, 1 mM EDTA, 1 mM EGTA, 1 mM DTT, Phosphatase Inhibitor Cocktail (Thermo Fisher Scientific, 78440), and the protease inhibitor cocktail (Roche, 11697498001). Samples were lysed by trituration through a 25-gauge needle 10 times on ice, rotated for 30 minutes at 4°C, and centrifuged (21000g, 30 minutes, 4°C) to obtain whole cell extracts. The protein concentration was determined by Bradford assay using protein assay dye (Bio-Rad). Lysates were boiled in 5x loading buffer (250 mM Tris–HCl, pH 6.8, 10% SDS, 0.05% bromophenol blue, 50% glycerol, 25% β −mercaptoethanol,) and then fractionated by SDS–PAGE. Western blot analysis was performed after electrophoretic separation of polypeptides by 4-15% SDS-PAGE (Bio-Rad, 5000006) and transfer to Immobilon-P/PVDF membrane (Millipore, IPVH00010). Blots were probed overnight at 4°C with the following primary antibodies: rabbit anti-vimentin at 1:2000 (Proteintech Group Inc, 10366-1-AP), rabbit anti-collagen I at 1:2000 (Proteintech Group Inc, 14695-1-AP), mouse anti-αSMA at 1:2000 (Abcam, ab7817) and rabbit anti-GAPDH at 1:5000 (GeneTex, GTX100118). Following incubation with primary antibodies, the blots were washed and then probed with the respective HRP-conjugated secondary antibodies at 1:5000 (Cell Signaling Technology, 7076 and 7074). Immuno-bands were subsequently detected by the enhanced chemiluminescence reaction (Thermo Fisher Scientific, 34095).

### Quantitative reserve transcription (RT) and real-time PCR

Total RNA from cell pellets and kidney tissues were isolated using TRIzol reagent (Invitrogen, 1559602) according to manufacturer’s instructions. cDNA was synthesized by Superscript II reverse transcriptase (Invitrogen, 18064071) with random hexamers. Primers for real-time PCR assays are listed in Supplementary Table S1. cDNA levels were determined quantitatively by real-time PCR using the Bio-Rad iTaq Universal SYBR Green Supermix system (1725125) and normalized with *Gapdh*. All results represent the mean ± SD of at least three independent experiments.

### Tissue preparation and histology

Kidneys were fixed in 4% paraformaldehyde (PFA) overnight at 4°C. For histological analysis, fixed tissues were embedded in paraffin wax, sectioned at 5 μm, and stained with hematoxylin and eosin (H&E) or trichrome staining by the Research Histology Facility at Yale School of Medicine. For immunofluorescence studies, fixed tissues were embedded in OCT (Sakura Finetek, 4583) and cryosectioned into 5-μm sections by Microscopy Core of the Department of Cellular and Molecular Physiology at Yale School of Medicine.

### Immunofluorescence

Cells seeded on slides were fixed in 4% PFA for 5 min. Slides with OCT-embedded tissue sections and cultured cells were washed with PBS twice and subsequently permeabilized with 0.5% Triton X-100/0.1% Tween-20/PBS at room temperature for 20 min. Slides were then blocked in blocking buffer (R.T.U. Animal Free Block and Diluent, Vector Laboratories, SP-5035-100) and incubated overnight with the following primary antibodies: rabbit anti-laminin (1:200, NB300-144, Novus Biologicals), chicken anti-LacZ (1:3000, BGL-1010, Aves Labs), rabbit anti-vimentin (1:200), mouse anti-α-SMA (1:200), rabbit anti-ARL13b (1:200, Proteintech Group, 17711-1-AP) and mouse anti-acetylated tubulin (1:5000, Sigma-Aldrich, clone 6-11B-1). Slides were washed three times in PBS and subsequently incubated for 1 h at room temperature in blocking buffer containing the secondary antibodies conjugated to Ig-Alexa Fluor 568, Ig-Alexa Fluor 488 or Ig-Fluorescein. Slides were washed three times in 1XPBS and then mounted with Vectashield Vibrance mounting medium with DAPI (Vector Laboratories, H-1800).

### Cyst index analysis

Cystic index was performed as previously described^29,41^. Cyst index was analyzed by Metamorph v.7.1 acquisition software (Universal Imaging) using H&E-stained images in sagittal sections of kidneys. Cystic index was calculated by dividing the cyst-containing area by the total kidney area.

### Blood Urea Nitrogen (BUN) measurement

Total blood was collected from mouse in a heparin-coated tube (BD Microtainer, BD Inc), and centrifuged (6000rpm, 5 minutes) to obtain plasma. Plasma BUN was measured using a Urea Nitrogen colorimetric detection kit (Invitrogen, EIABUN) following the manufacturer’s instructions.

### Statistical analysis

Data are presented as mean with error bars indicating the standard deviation (S.D.). Statistical significance was calculated by unpaired two-tailed t test and Pearson correlation analysis using GraphPad Prism 8.

## Disclosures

None.

## Funding

This work was supported by National Institute of Health grants R01DK113135 and R01HD093608 (to Dr. Sun).

## Acknowledgements

We thank the Liem lab for *Gli1^lacZ^* mice; the members of Somlo laboratory and Sun laboratory for helpful discussions; A. Cox for critical reading of the manuscript; S. Mentone in the Microscopy and Imaging Core of Cellular and Molecular Physiology Department of Yale School of Medicine for histology assistance.

---

**Supplementary Table S1:**
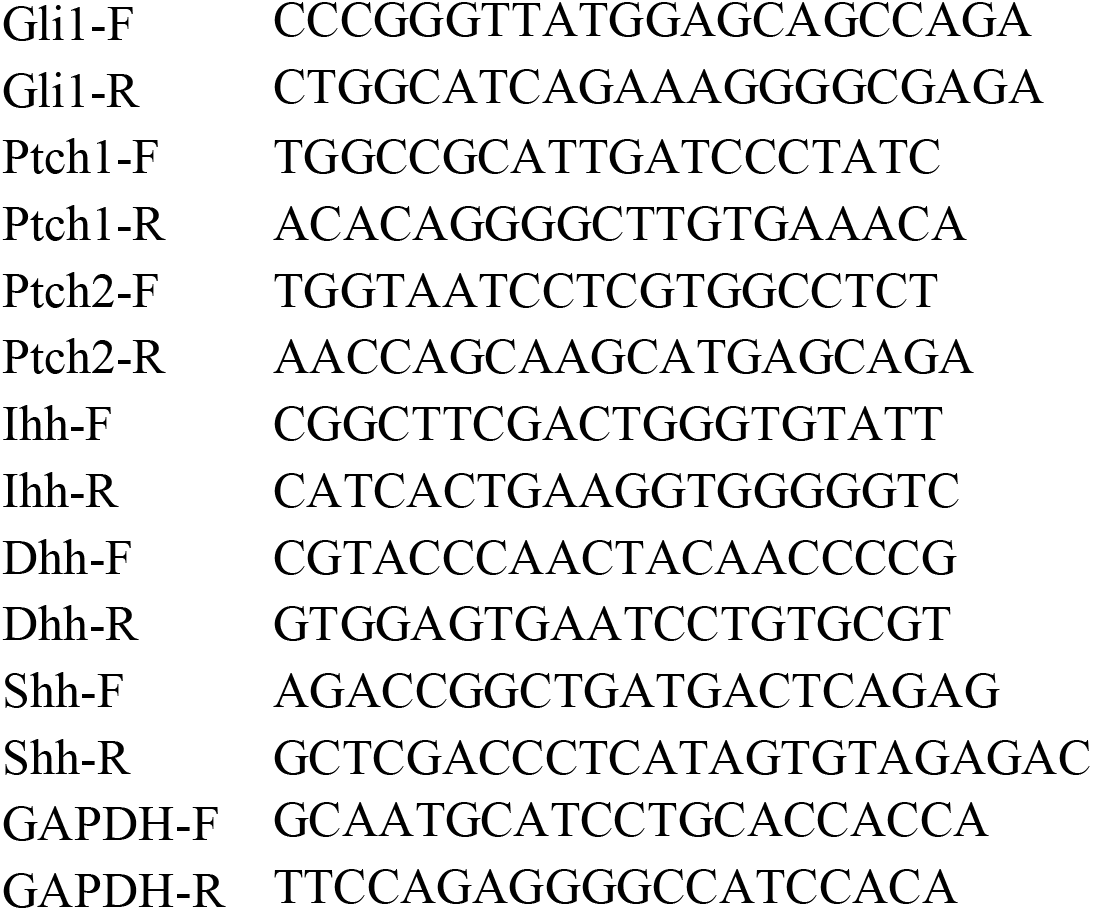
Primers used in qRT-PCR:

## Notes

### Competing Interest Statement

The authors have declared no competing interest.

